# Comparison Between Non-Catheterized And Catheter Associated Urinary Tract Infections Caused By Extended Spectrum B-Lactamase Producing *Escherichia Coli* And *Klebsiella Pneumoniae*

**DOI:** 10.1101/019026

**Authors:** Fouzia Ashraf, Shagufta Iram, Gul-e-zar Riaz, Farhan Rasheed, Mahmood Shaukat

## Abstract

Catheter associated urinary tract infection’s (CAUTI) account for more than 80% of all healthcare associated infections (HAIs) as compared to non-catheterized urinary tract infections. Catheter associated urinary tract infections occur on the third day after insertion of catheter in patients having urinary tract infection (UTI). In long term catheter use, bio-film form along the catheter which increases the risk of antibiotic resistant pathogens. Most common pathogens involved were Escherichia coli and Klebsiella pneumoniae which produce the enzymes Extended Spectrum β-Lactamases (ESBLs).

**Objective:** To compare the frequency of ESBLs in catheterized and non-catheterized UTI infections.

**Materials and Methods:** This comparative study was conducted at the Microbiology Department, Allama Iqbal Medical College, Lahore, from June 2014 to January 2015. Urine samples were cultured according to WHO protocol and antimicrobial Susceptibility testing was performed by *Modified Kirby-Bauer disc diffusion Method.* Escherichia coli and Klebsiella pneumoniae were tested for ESBL production by phenotypic confirmatory method of disk diffusion synergy using a disc of amoxicillin-clavulonate (30μg) and ceftrixone (30μg), cefotaxime (30μg) and aztreonam (30μg) discs.

**Results:** Out of 300 positive urinary isolates of Escherichia coli and Klebsiella pneumoniae from CAUTI, 65.33% were ESBL producing isolates whereas out of 300 positive urinary isolates of Escherichia coli and Klebsiella pneumoniae from non-catheterized UTI, 47.66% were ESBL producing isolates. The results were highly significant (p < 0.001).

**Conclusion:** Results showed that frequency of ESBLs were higher in catheterized patients as compared to non-catheterized patients. This is suggestive of a need for regular screening and surveillance for ESBL producing organisms. Patients infected with these organisms should be nursed with contact precautions to avoid the spread of nosocomial infection.

## Introduction

Infections that patients acquire during hospital stay and receiving treatment for medical and surgical conditions are healthcare-associated infections [1]. Urinary tract infection is the most common bacterial infection that occurs in urinary tract and accounts for 40% of all hospital acquired infections [2]. The most common healthcare-associated infection is catheter-associated urinary tract infections which occur in patients after insertion of catheter and accounts for 30% to 40% of all the hospital acquired infections [3, 4]. Insertion of catheter in patients greater than 7 days accounts infection up to 25% and increases to 100% after 30 days [4]. Bacterial colonies form into the epithelial cells of the urinary tract which are responsible for the infection [5]. The infection that occurs with an indwelling Foley’s urinary catheter within 48 hours is a catheter-associated urinary tract infection [6]. Indwelling Foley’s urinary catheter enables emptying of the bladder. Urinary catheterization is categorized as short term that is in situ less than 28 days or long term is in situ more than 28 days [7]. The presence of bacteria in urine is known as bacteriuria which is either asymptomatic (CA-ASB) or symptomatic bacteriuria. Asymptomatic bacteriuria is usually present in CAUTI [8]. Patients with urinary tract infections have symptoms of fever, urinary urgency, lower abdominal pain, dysuria, pyuria, and leukocytes in urine [9]. The presence of bacteria with a count of 10^5^ CFUs per ml is an indication of urinary tract infection [10]. The level of bacteria in urine increases from 10^5^ to 10^8^ CFU per ml within 24 to 48 hours if antimicrobial therapy is not given to the patient [11]. Bio-film formation along the catheter reflects the presence of organisms that increase resistance to the host immune response and to antibiotics. Bio-film forms when a catheter was inserted greater than 2 weeks ago [12]. Most common pathogens associated with CAUTI are *Escherichia coli, Klebsiella pneumoniae, Enterobacter species, Proteus species, Pseudomonas aeruginosa, S. saprophyticus, Candida species.* Among gram negative rods, *Escherichia coli* and *Klebsiella pneumoniae* are more prone to produce ESBLs [2]. Extended spectrum β-lactamases (ESBLs) are enzymes that are often located on plasmids. These plasmid mediated β-lactamases are capable of hydrolyzing and inactivating extended spectrum β-lactams with an oxyimino side chain like cephalosporins (cefotaxime, ceftriaxone, and ceftazidime) and oxyimino-monobactam (aztreonam) [13]. Previous use of antibiotics, catheterization, prolonged hospital stay, and invasive infections are the major risk factors for ESBL production [14]. Morbidity and mortality of ESBLs increased in both developed and developing countries. Therefore the present study was planned to isolate ESBLs.

## Materials and Methods

### Study population

This study was conducted at the Microbiology Department of AIMC, Lahore. From June 2014 to January 2015, consecutive, non-duplicate isolates of *Escherichia coli* and *Klebsiella pneumoniae* were collected from urine samples of patients admitted in different wards of Jinnah Hospital who were hospitalized for at least 48 hours. Catheterized patients having UTI’s with underlying diseases such as spinal cord injuries, renal diseases, chronic UTI’s, sepsis, diabetes, hepatitis, and urological operations belonging to the indoor department were included in this study. Ages ranged from 2 to 85 years of either gender. Non-catheterized patients having UTI with symptoms of fever, cost-vertebral angle pain, urinary frequency, burning micturition, bacteriuria, leukocytes and pyuria in urine regardless of age and sex were included. Bacterial colonies of either *Escherichia coli* or *Klebsiella pneumoniae* on culture plates with 10^5^ CFUs per ml was considered as UTI diagnosis. A positive double disc diffusion test was considered for ESBLs detection. Patients of less than 2 years were excluded. Informed consent was taken from either the patient or a relative.

### Sample Collection

Two ml urine samples were collected from catheterized patients. In patients with short term catheterization, samples were obtained by puncturing the catheter tubing with a needle or syringe using aseptic techniques. In patients with long term indwelling catheters, urine samples were collected from the freshly placed catheter. In symptomatic patients, urine samples were collected immediately prior to antimicrobial therapy. Mid-stream urine samples were collected in a sterile urine container from non-catheterized patients.

### Urine Culturing

The urine specimens were subjected to urine culture on Cysteine Lactose Electrolyte Deficient Agar (CLED) and streaked for colony count. The plates were incubated at 37°C for 16 to 18 hrs and then examined for bacterial growth. Organisms present on culture were identified by standard microbiological procedures (Colonial morphology, Gram’s stain appearance, motility, routine biochemical tests). Antimicrobial susceptibility testing was performed by Modified Kirby Bauer disc diffusion method on Mueller-Hinton agar. In this method, the inoculums were adjusted to 0.5 McFarland standards and swabbed onto the surface of Mueller-Hinton agar with the help of a swab. The antibiotic discs were placed onto the agar plate and incubated at 37°C for 16-18 hours. After incubation, zones of inhibition were noted and compared with reference zones. The following antibiotics discs were used: cefotaxime (30μg) / cefpodoxime (30μg) / ceftriaxone (30μg) / ceftazidime (30μg) / aztereonam (30μg) and amoxicillin-clavulanate (30μg) for the ESBL producing *Escherichia coli* and *Klebsiella pneumoniae* on Mueller-Hinton agar. These ESBL producing organisms showed a zone diameter of ≤ 17 mm for cefpodoxime / ≤ 22 mm for ceftazidime / ≤ 27 mm for aztreonam / ≤ 25 mm for ceftriaxone or ≤ 27 mm for cefotaxime as recommended by CLSI (Clinical and Laboratory Standards Institute) [15]. ESBL detection was confirmed by double disc diffusion synergy method. In this method, a synergy was determined between a disc of amoxicillin-clavulanate (30μg) and a 30μg disc of each third generation cephalosporin test antibiotic (ceftrixone and cefpodoxime) placed at a distance of 25-30 mm from center to center on a Mueller-Hinton Agar, (MHA) plate swabbed with the isolates of *Escherichia coli* or *Klebsiella pneumoniae.* The plates were incubated at 37°C for 16-18 hours. After incubation, augmentation or key-hole phenomenon was observed which indicated the production of ESBL [16]. *Escherichia coli* American Type Culture Collection (ATCC^®^) 25922 and *Klebsiella pneumoniae* (ATCC^®^) 700603 were used as control.

### Statistical Analysis

SPSS version-20 was used for statistical data analysis. Frequency tables and percentages expressed as categorical variables and continuous variables expressed as mean and standard deviation. In statistical analysis, the value of p less than 0.05 is considered significant and a value of p greater than 0.05 is considered insignificant. Tests of significance were calculated using chi-square test for ESBL positivity. The p-value of less than 0.05 was considered statistically significant.

## Results

300 isolates (*Escherichia coli* and *Klebsiella pneumoniae*) were included from catheterized and non-catheterized patients from a total of 1012 urine specimens received during the study’s duration. From 1012 urine samples, 506 urine samples were from each of catheterized patients and non-catheterized patients. Out of a total of 506 samples from catheterized patients, 54 samples did not show bacterial growth while 452 had positive cultures. In non-catheterized patients, out of 506 samples, 67 samples did not show bacterial growth while 439 had positive cultures. Out of 452 positive cultures from catheterized patients, 152 were urinary pathogens *i.e Staphylococcus Saprophyticus, Proteus vulgaris, Pseudomonas aeroginosa, Candida* species and 300 were *Escherichia coli* and *Klebsiella pneumoniae* while in non-catheterized patients, out of 439 positive cultures, 139 were other urinary pathogens and 300 were *Escherichia coli* and *Klebsiella pneumoniae*.

Out of 300 *Escherichia coli* and *Klebsiella pneumoniae* from catheterized patients, 56% (168) were *Escherichia coli* and 44% (132) were *Klebsiella pneumoniae* where as in non-catheterized patients, out of 300 *Escherichia coli* and *Klebsiella pneumoniae*, 50.7% (152) were *Escherichia coli* and 49.3% (148) were *Klebsiella pneumoniae.* Out of 56% (168) *Escherichia coli* and 44% (132) *Klebsiella pneumoniae* from catheterized patients, 75% (126) were ESBL producing *Escherichia coli* and 53.03% (70) were ESBL producing *Klebsiella pneumoniae.* Where as in non-catheterized patients out of 50.7% (152) *Escherichia coli*, 71.05% (108) were ESBL producing *Escherichia coli* and out of 49.3% (148) *Klebsiella pneumoniae*, 23.64% (35) were ESBL producing *Klebsiella pneumoniae* as shown in table 1.

**Table 1:** Bacterial growth in patients of urinary tract infection

| Bacterial growth | Catheterized patients having UTI | Non-catheterized patients having UTI | p-value |
| --- | --- | --- | --- |
| *Escherichia coli* | 168 (56%) | 152 (50.7%) | 0.190* |
| *Klebsiella pneumoniae* | 132 (44%) | 148 (49.3%) |  |
| Total | 300 (100%) | 300 (100%) |  |

\* P > 0.05 (not significant)

Therefore, a total of 65.33% (196) isolates were positive for ESBLs from catheterized patients while in non-catheterized patients, a total of 47.66% (143) isolates were positive for ESBLs. The results of ESBLs from catheterized and non-catheterized UTI patients were highly significant (p= 0.001). Double disc synergy method positive for ESBLs as shown in the figure 1:

**Figure 1:**
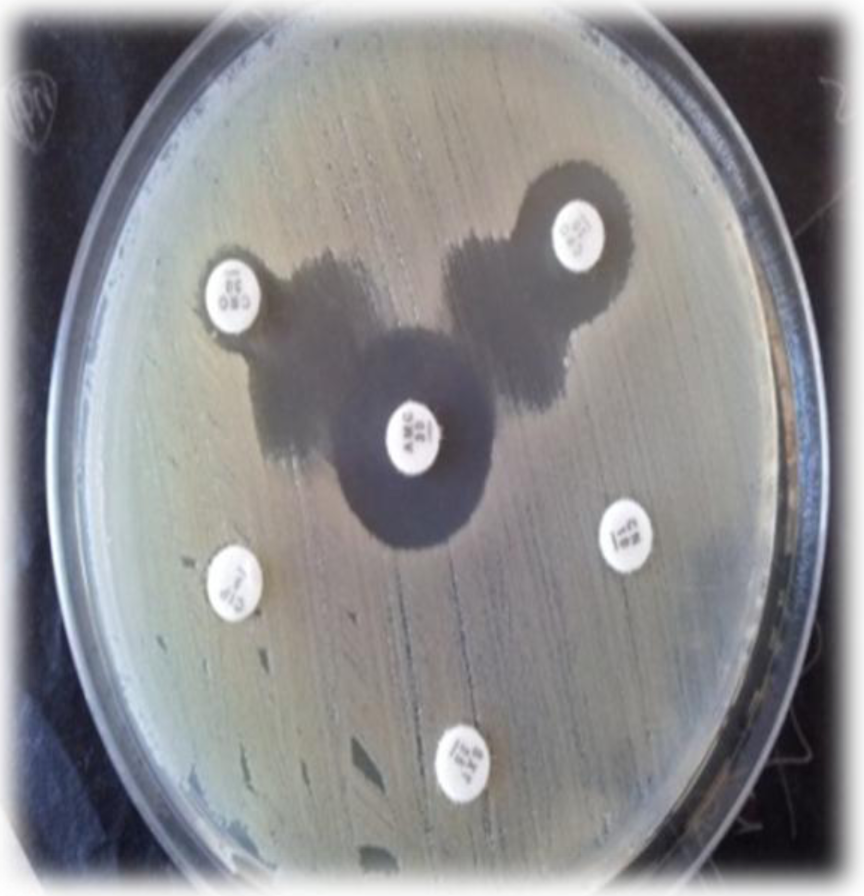
Double disc synergy test for ESBL producing organism.

Mean age of patients with a urinary tract infection was 39.13 ± 19.582 (minimum age 2 and maximum age 85). Mean age of patients with catheter associated urinary tract infection was 39.61 ± 19.289 and mean age of non-catheterized UTI patients was 38.65 ± 19.891. There was no statistically significant difference between catheterized and non-catheterized UTI patients (p=0.878). Patients of 2-12 years with catheter associated UTI were 5.7% (17) where as non-catheterized UTI were 6.7% (20). Patients of 13-30 years with CAUTI were 35% (105) where as non-catheterized UTI were 36.3% (109). Patients of 31-60 years with CAUTI were 44% (132) while non-catheterized UTI were 41% (123). Patients of 61 years and above with catheter associated UTI were 15.3% (46) where as non-catheterized UTI were 16% (48) as shown in table 2.

**Table 2:** Age distribution among Catheterized patients and Non-catheterized patients having UTI

| Age groups | Catheterized patients having UTI | Non-catheterized patients having UTI | p-value |
| --- | --- | --- | --- |
| 2-12 years | 17 (5.7%) | 20 (6.7%) | 0.878* |
| 13-30 years | 105 (35%) | 109 (36.3) |  |
| 31-60 years | 132 (44%) | 123 (41%) |  |
| 61 years and above | 46 (15.3%) | 48 (16%) |  |
| Total | 300 (100%) | 300 (100%) |  |

\* $p > 0.05$ (not significant)

From catheterized UTI patients, 60.3% (180) were male and 39.7% (119) were female whereas from non-catheterized UTI patients, 54.3% (163) were male and 45.7% (137) were female. There was no statistical difference between the two groups (p= 0.137).

Catheterized UTI patients belonged to the Urology Ward were 6.3% (19), Medicine 35.3% (106), ICU 19% (57), Paeds 4.3% (13), Surgery 16.7% (50), Orthopeadic 9.7% (29), Pulmonology 5.7% (17), Gynecology 2.7% (8), and Oncology 0.3% (1). Non-catheterized UTI patients belonged to Urology Ward were 8.7% (26), Medicine 43.3% (130), ICU 20.3% (61), Paeds 5.7% (17), Surgery 10.3% (31), Orthopaedic 2% (6), Pulmonology 7.3% (22) and Gynecology 2.3% (7) and are shown in figure 2. The number of patients belonging to different wards were statistically significant (p = 0.001). 45.7% (137) of the catheterized patients who had UTI’s used previous antibiotics of cephalosporin’s and 37.7% (113) of non-catheterized patients used cephalosporin antibiotics that were statistically significant (p = 0.047).

**Figure 2:**
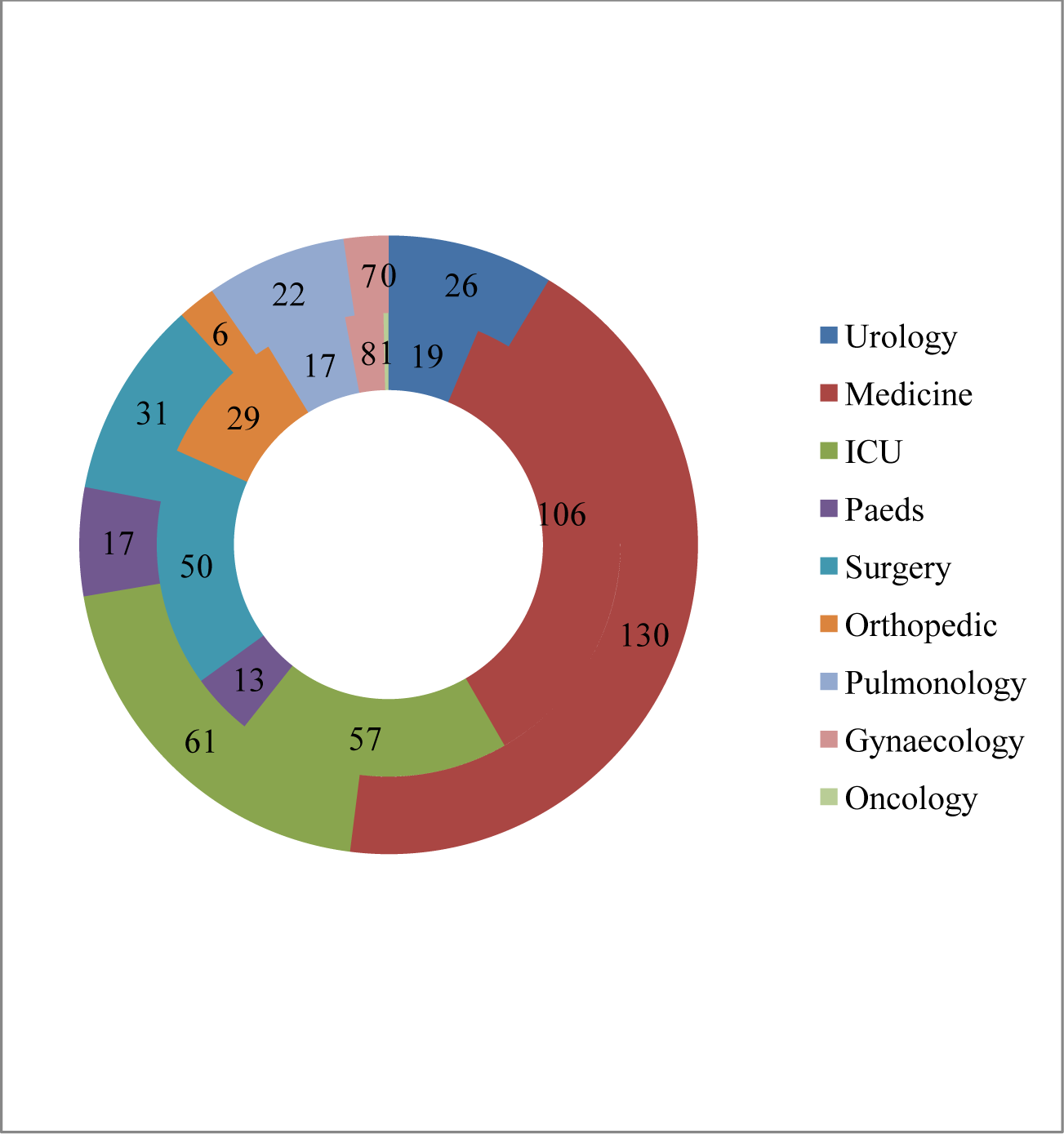
Smaller ring shows number of catheterized patients and larger ring shows number of non-catheterized patients in different wards.

## Discussion

Extended spectrum β-lactamases production was identified in the early 1980s. *Escherichia coli* and *Klebsiella pneumoniae* were the predominant ESBLs producing organisms [17, 18]. Extended spectrum β-lactamases produced by most common urinary pathogens are increasing in number. They are resistant to β-lactams that are difficult to treat by clinicians. Inappropriate use of antibiotics led to the increase in resistance to the antibiotics that make the treatment difficult. In this study 45.7% of catheterized patients and 37.7% non-catheterized patients have previously used the antibiotics cephalosporin which leads to the failure of antimicrobial therapy due to resistance of *Escherichia coli* and *Klebsiella pneumoniae* as shown in Table 3. Taneja *et al* (2008) reported the highly drug resistant uropathogens for testing ESBL production and 36.5% were found to be ESBL producers. The high sensitivity was found in *Klebsiella pneumoniae* as compared to *Escherichia coli* [19]. These percentages are similar to our data. Prior antibiotic use was the risk factor for ESBL-producing *Escherichia coli* or *Klebsiella pneumoniae* infection [20]. Isolates associated with CAUTI also form biofilm which enables them to survive against antimicrobials. In a long term catheterized patient, bio-film forms along the catheter which causes the micro-organisms to adhere to it.

**Table 3:** Previous antibiotic use of patients

| Antibiotic use | Catheter associated urinary tract infection | Non-catheter associated urinary tract infection | p -value |
| --- | --- | --- | --- |
| Patients with previous antibiotic use | 137 (45.7%) | 113 (37.7%) | 0.047* |
| Patients with no previous antibiotic use | 163 (54.3%) | 187 (62.3%) |  |
| Total | 300 (100%) | 300 (100%) |  |

\*P < 0.05 (significant)

These cause persistent infections, survive for a long duration, and finally become resistant to antimicrobial therapy [21]. In our study, the frequency of ESBLs was 65.33% from catheterized patients and 47.66% from non-catheterized patients as shown in Table 4. Afridi *et al* (2011) from Pakistan, reported a prevalence of 65.7% ESBL producing organisms from urine of non-catheterized patients [22]. Another study from Nagpur, India showed the prevalence of ESBLs to be 48.3% in urinary isolates [23].

**Table 4:** ESBLs in patients of urinary tract infection

| ESBLs production | Catheterized patients having UTI | Non-catheterized patients having UTI | p-value |
| --- | --- | --- | --- |
| ESBLs | 196 (65.3%) | 143 (47.7%) | 0.001* |
| Non-ESBLs | 104 (34.7%) | 157 (52.3%) |  |
| Total | 300 (100%) | 300 (100%) |  |

\*p < 0.05 (highly significant)

ESBL producing *Escherichia coli* showed the highest frequency (75%, 71.05%) followed by *Klebsiella pneumoniae* (53.03%, 23.64%) in both catheterized and non-catheterized patients positive for UTI’s. This presented evidence that treatment with beta-lactam antibiotics is the risk factor of acquiring ESBL producing *Escherichia coli* in CAUTI. Spadafino *et al* (2014) reported 83.4% ESBL producing *Escherichia coli* in CAUTI [24]. Y.S. Yang *et al* (2010) reported 58.3% ESBL producing *Escherichia coli* and 41.7% *Klebsiella pneumoniae* in urinary tract infections [17]. Hafeez *et al* (2009) reported 44.8% *Escherichia coli* and 38.6% *Klebsiella pneumoniae* among urinary pathogen isolates [25]. The mean age of patients was 39.13 years. A study from Pakistan reported a mean age of 47 years [26]. The frequency of male patients was 60.3% in catheterized patients and 54.3% in non-catheterized patients. 39.7% were female catheterized patients and 45.7% were in non-catheterized patients. There was no statistical difference between male and female patients (p = 0.137). The highest number of patients belonged to the Medicine Ward. In our region, there is a strong positive association between prior antibiotic use and ESBL production of the catheterized and non-catheterized patients (p = 0.047). 45.7% of catheterized patients and 37.7% of non-catheterized patients have previously used 3^rd^ generation cephalosporins. The urine samples of these patients showed resistance of *Escherichia coli* and *Klebsiella pneumoniae* to the beta lactam drugs and are a major risk factor for ESBLs production. The only choice of drug for ESBL producing organisms is imipenem. The duration of catheterization and misuse of antibiotics were also risk factors for the emergence of resistance to drugs and thus treatment of these patients becomes difficult. It is necessary that catheterization in situ should not be greater than 3 days. In-out catheterization pattern should follow hygienic conditions to decrease the spread of nosocomial infections and ESBL production.

## Conclusion

A high frequency of ESBL producing organisms was found especially among *Klebsiella pneumoniae* and *Escherichia coli* in urinary isolates. ESBL producing organisms were found higher amongst catheterized patients associated with urinary tract infections as compared to non-catheterized patients associated with urinary tract infections. This is suggestive of a need for regular screening and surveillance for ESBL producing organisms. Since ESBLs destroy the cephalosporins given as first line antibiotics in hospitals, this leads to increased morbidity and mortality and therefore ESBLs are clinically important. Thus, accurate detection of these organisms is essential for effective treatment. ESBL production from clinical specimens also reflects non-judicious use of antimicrobials. Patients infected with these organisms should be nursed with contact precautions to avoid the spread of nosocomial infection. Further typing and sub-typing of ESBLs based on gene sequencing by PCR is planned as continuation of this study. This will show the most common ESBL types in our clinical settings. This confers antibiotic resistance to specific group of antibiotics. It will equip the treating physicians with better treatment options for patients infected with ESBL producing organisms.

## Acknowledgements

**Shahzeb Javed**: Medical Student at King Edward Medical University. Helped prepare manuscript and assisted with samples collection.

## References

Hughes, R.G. and A.S. Collins, Preventing health care– associated infections. 2008.

Patil, A.B., et al., Catheter-associated urinary tract infection: Aetiology, ESBL production, and risk factors. Journal, Indian Academy of Clinical Medicine, 2014. 15(1): p. 23.

Conway, L.J. and E.L. Larson, Guidelines to prevent catheter-associated urinary tract infection: 1980 to 2010. Heart & Lung: The Journal of Acute and Critical Care, 2012. 41(3): p. 271&283.

Jaggi, N. and P. Sissodia, Multimodal Supervision Programme to Reduce Catheter Associated Urinary Tract Infections and Its Analysis to Enable Focus on Labour and Cost Effective Infection Control Measures in a Tertiary Care Hospital in India. Journal of Clinical and Diagnostic Research : JCDR, 2012. 6(8): p. 1372&1376.

Sheerin, N.S., Urinary tract infection. Medicine, 2011. 39(7): p. 384&389.

Conway, L.J. and E.L. Larson, Guidelines to prevent catheter-associated urinary tract infection: 1980 to 2010. Heart & lung : the journal of critical care, 2012. 41(3): p. 271&283.

Guidelines for the prevention of catheter-associated urinary tract infection. 2011, Dublin: HSE Health Protection Surveillance Centre on behalf of SARI.

Gould, C.V., et al., Guideline for prevention of catheter-associated urinary tract infections 2009. Infection Control, 2010. 31(04): p. 319&320.

Hooton, T.M., et al., Diagnosis, Prevention, and Treatment of Catheter-Associated Urinary Tract Infection in Adults: 2009 International Clinical Practice Guidelines from the Infectious Diseases Society of America. Clinical Infectious Diseases, 2010. 50(5): p. 625&663.

Schmiemann, G., et al., The Diagnosis of Urinary Tract Infection: A Systematic Review. Deutsches Ärzteblatt International, 2010. 107(21): p. 361&367.

Tambyah, P.A. and D.G. Maki, Catheter-associated urinary tract infection is rarely symptomatic: a prospective study of 1497 catheterized patients. Archives of internal medicine, 2000. 160(5): p. 678&682.

Nicolle, L.E., Catheter associated urinary tract infections. Critical Care, 2014. 1: p. 4.1.

Rupp, M.E. and P.D. Fey, Extended spectrum β-lactamase (ESBL)-producing Enterobacteriaceae. Drugs, 2003. 63(4): p. 353&365.

Kurtaran, B., et al., Antibiotic resistance in community-acquired urinary tract infections: prevalence and risk factors. Medical Science Monitor, 2010. 16(5): p. CR246&CR251.

Cockerill, F.R., Performance standards for antimicrobial susceptibility testing: twenty-first informational supplement. 2011: Clinical and Laboratory Standards Institute (CLSI).

Paterson, D.L. and R.A. Bonomo, Extended-Spectrum β-Lactamases: a Clinical Update. Clinical Microbiology Reviews, 2005. 18(4): p. 657&686.

Yang, Y.-S., et al., Impact of extended-spectrum β-lactamase-producing Escherichia coli and Klebsiella pneumoniae on the outcome of community-onset bacteremic urinary tract infections. Journal of Microbiology, Immunology and Infection, 2010. 43(3): p. 194&199.

Jacoby, G.A. and A.A. Medeiros, More extended-spectrum beta-lactamases. Antimicrobial Agents and Chemotherapy, 1991. 35(9): p. 1697.

Taneja, N., et al., Occurrence of ESBL & Amp-C beta-lactamases & susceptibility to newer antimicrobial agents in complicated UTI. The Indian journal of medical research, 2008. 127(1): p. 85&88.

Lautenbach, E., et al., Extended-spectrum β-lactamase-producing Escherichia coli and Klebsiella pneumoniae: risk factors for infection and impact of resistance on outcomes. Clinical Infectious Diseases, 2001. 32(8): p. 1162&1171.

Trautner, B.W. and R.O. Darouiche, Role of biofilm in catheter-associated urinary tract infection. American journal of infection control, 2004. 32(3): p. 177&183.

Afridi, F.I., B.J. Farooqi, and A. Hussain, Frequency of extended spectrum beta lactamase producing enterobacteriaceae among urinary pathogen isolates. J Coll Physicians Surg Pak, 2011. 21: p. 741-744.

Tankhiwale, S.S., et al., Evaluation of extended spectrum beta lactamase in urinary isolates. Indian J Med Res, 2004. 120(6): p. 553&6.

Spadafino, J.T., et al., Temporal trends and risk factors for extended-spectrum beta-lactamase-producing Escherichia coli in adults with catheter-associated urinary tract infections. Antimicrobial Resistance and Infection Control, 2014. 3(1): p. 599.

Hafeez, R., et al., Frequency of extended spectrum beta lactamase producing gram negative bacilli among clinical isolates. Biomedica, 2009. 25(2): p. 112&5.

Jabeen, K. and R. Hasan, Frequency and sensitivity pattern of extended spectrum beta lactamase producing isolates in a tertiary care hospital laboratory of Pakistan.

